# Prediction of Electric Fields Induced by Transcranial Magnetic Stimulation in the Brain using a Deep Encoder-Decoder Convolutional Neural Network

**DOI:** 10.1101/2022.10.27.513583

**Authors:** Mohannad Tashli, Muhammad Sabbir Alam, Jiaying Gong, Connor Lewis, Carrie L. Peterson, Hoda Eldardiry, Jayasimha Atulasimha, Ravi L. Hadimani

**Author notes:** Equal contributions, author order arbitrary (Mohannad Tashli, Muhammad Sabbir Alam, Jiaying Gong).

## Abstract

Transcranial magnetic stimulation (TMS) is a non-invasive, effective, and safe neuromodulation technique to diagnose and treat neurological and psychiatric disorders. However, the complexity and heterogeneity of the brain composition and structure pose a challenge in accurately determining whether critical brain regions have received the right level of induced electric field. Numerical computation methods, like finite element analysis (FEA), can be used to estimate electric field distribution. However, these methods need exceedingly high computational resources and are time-consuming. In this work, we developed a deep convolutional neural network (DCNN) encoder-decoder model to predict induced electric fields, in real-time, from T1-weighted and T2-weighted magnetic resonance imaging (MRI) based anatomical slices. We recruited 11 healthy subjects and applied TMS to the primary motor cortex to measure resting motor thresholds. Head models were developed from MRIs of the subjects using the SimNIBS pipeline. Head model overall size was scaled to 20 new size scales for each subject to form a total of 231 head models. Scaling was done to increase the number of input data representing different head model sizes. Sim4Life, a FEA software, was used to compute the induced electric fields, which served as the DCNN training data. For the trained network, the peak signal to noise ratios of the training and testing data were 32.83dB and 28.01dB, respectively. The key contribution of our model is the ability to predict the induced electric fields in real-time and thereby accurately and efficiently predict the TMS strength needed in targeted brain regions.

## I. Introduction

**T**RANSCRANIAL magnetic stimulation (TMS) is a safe, effective, and non-invasive treatment for several psychiatric and neurological disorders [1], [2]. TMS is FDA approved and widely used for the treatment of major depressive disorder and obsessive-compulsive disorder [3]–[5]. Reports on the development of improved TMS devices for the treatment of brain diseases and disorders have recently increased [6]–[13].

The majority of these techniques use an electromagnetic signal targeting specific brain regions to treat brain disorders. To prevent undesired or excessive brain stimulation, the electromagnetic pulses must target specific neuronal subpopulations in the brain with a specific threshold magnetic field amplitude and induce an electric field (E-field) of approximately 150 V/m depending on the type of neurons targeted [14]–[17].

Normal anatomical diversity and morphology of the human brain cause considerable variations in the effective TMS location, intensity, and E-Field distribution [18]. These characteristics might account for significant variations reported in both diagnostic parameters and clinical outcomes in terms of efficacy and remission [19]. Therefore, the TMS intensity applied to initiate action potentials in motor cortex neurons varies across subjects in part due to varying brain anatomy [20]. The threshold E-field can be identified by measuring motor evoked potential (MEP) in the first dorsal interosseous muscle in the thumb. When the MEP is 50μV or more in five out of ten consecutive stimuli, the corresponding stimulator output power is considered the resting motor threshold (RMT) [21], [22]. RMT is obtained in a subject when the E-field reaches a threshold field of ≈150 V/m in the target area of the brain. The threshold E-field is calculated using finite element analysis of the subject’s head model generated from magnetic resonance images (MRIs). Measuring RMT or calculating the induced E-field requires a significant amount of time and computational resources [23]–[26]. Also, the induced E-field calculated from the MRI does not consider the brain’s functional connectivity and resting functional state. However, for many clinical purposes, predicting the induced E-field will allow clinicians and researchers to better understand the relationship between the TMS dosage (as a percent of RMT) and neuroanatomy. In this study, we predicted the induced E-field distribution in brain models using simulated TMS and machine learning algorithms to overcome the limitations of finite element analysis.

E-fields can be predicted by developing a learning-based method that utilizes current advancements in deep convolutional neural networks (DCNN). DCNNs are completely trainable, multi-layered models that can recognize and depict intricate, high-dimensional input-output correlations [27], [28]. These techniques are widely used in several areas of computer vision and problems related to image segmentation [29]–[31]. DCNNs can be useful in reducing simulation time for prediction-related tasks. Hence, we developed a DCNN-based encoder-decoder network to predict induced E-fields from MRI-based anatomical images for a given (fixed) position and orientation of the TMS coil.

A few studies have considered using artificial neural networks for predicting E-fields during TMS [32]–[34]. With the DCNN model, once trained, the induced E-field estimation requires less time and less computational resources compared to FEA-based methods. In addition, subjects requiring TMS will not be exposed to unnecessary stimulation as the DCNN model can predict the TMS responses for the specific subject. Other studies [35], [36] slightly modified the DCNN algorithms to overcome a few limitations and achieved a better performance in predicting E-fields compared to previous studies. Previous studies have relied on a public database to obtain MRIs and augment the data by rotating the coil position, which does not represent the clinical conditions as the coil position and rotation do not change significantly compared to the shape and size of the head.

The purpose of this study is to accurately predict the induced E-field on the subjects’ head models using anatomical and E-field data collected through our methodological processes. Furthermore, simulations were performed for therapeutic purposes where the TMS coil position is specific with respect to the subject’s anatomy. In this work, we have scaled each head model and generated 20 new head models’ scales for each subject to represent variation in neuroanatomy in a clinical setting.

## II. Methodology

Eleven healthy individuals (seven females and four males, 24.6 ± 5 years) participated in this study. All participants were screened to ensure the safety of the TMS and MRI protocols and provided informed consent. This study was approved by the Virginia Commonwealth University Institutional Review Board.

### A. TMS Experiments in Human Subjects

Each participant completed one TMS session targeting the first dorsal interosseous, described in detail in Mittal et al. [20]. Single pulse TMS was delivered as a monophasic posterior– anterior current to the primary motor cortex contralateral to the resting arm using a Magstim BiStim^2^ stimulator via a 70-mm figure-of-eight coil (P/N 4150-00). The hotspot for the target muscle was identified as the location evoking the largest peak-to-peak amplitude MEP using the lowest stimulation intensity [37], [38]. RMT was defined as the lowest stimulus intensity that induced MEPs of ≥50 μV in at least 5 of 10 consecutive stimuli with the target muscle fully relaxed [39].

### B. Head Models

Eleven healthy individuals were recruited, and MRI scanned as described in Mittal et al [20], [40] by a Phillips 3.0T system. Example structural T1 and T2 weighted MRI scans are shown in Fig. 1. Using the extracted T1- and T2-weighted images, a SimNIBS pipeline (SimNIBS Developers 2019, v2.0.1) [19], [23] was used to create head models (Fig. 2) consisting of seven separate segments (white matter, gray matter, cerebrospinal fluid, skin, skull, ventricles, and cerebellum) as separate 3D modeled files. Abnormalities were smoothed using Meshmixer (AutoDesk, Inc. v11.2.37) [40].

**Fig. 1.**
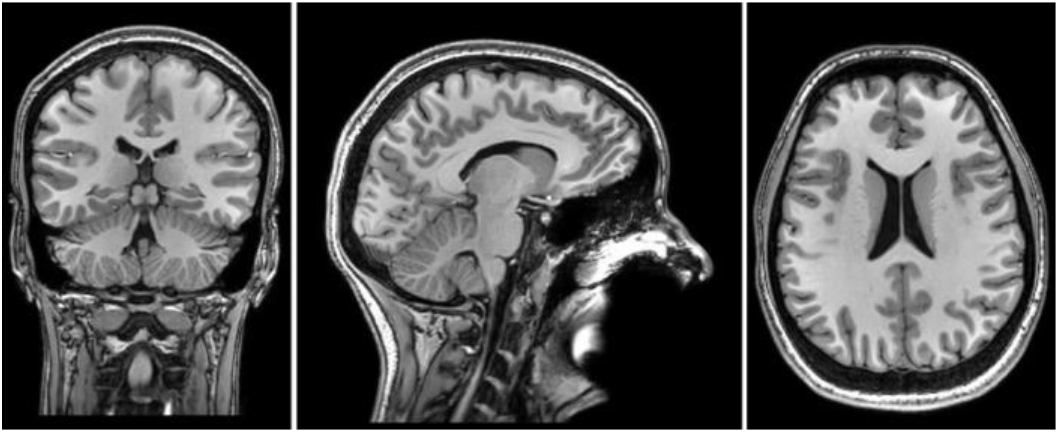
A study participant original MRI coronal, sagittal and horizontal views.

**Fig. 2.**
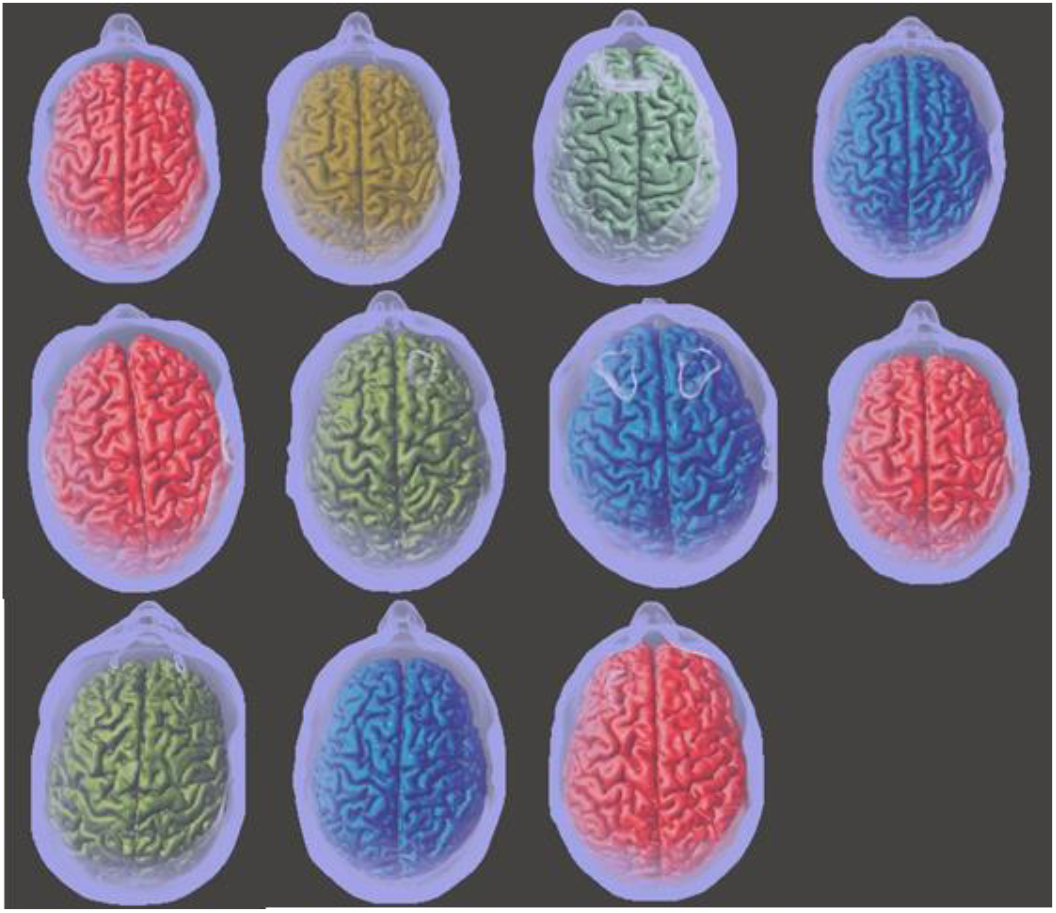
Head model variations in 11 subject participants before scaling.

### C. Head Models Scaling

The original head model’s segments’ sizes went through an overall size scaling to generate more input data for machine learning algorithms. The scaling process was equally performed on all three dimensions using Meshmixer (AutoDesk, Inc. v11.2.37). Each head model was scaled to 20 distinct scales, ranging from a scale of 0.9 to 1.1 with an increment of 0.01. That yields 21 head models for each subject including the original scale, which gave us 231 distinct head models.

### D. TMS Simulation

Sim4Life finite element analysis software (Zurich Med Tech, v6.2.1.4972) was used to compute induced E-field from peak intensity stimulation of the primary motor cortex from the head models [17], [23], [25], [41]. The simulated coil and positioning matched the empirical setup, targeting the precentral gyrus posterior to the superior frontal sulcus within the “knob” as defined by Yousry et al. [40], [42]. The coil was fixed at this position for all simulations performed, as shown in Fig. 3. Simulations were carried out on 231 head models with an amplitude of 5000 Amps at a stimulation frequency of 2.5 kHz. Material properties of the individualized segments and surrounding air were selected from the IT’IS LF database (IT’IS Foundation, v4.0) [40] for skin, skull, grey matter, white matter, cerebrospinal fluid, ventricles, and cerebellum.

**Fig. 3.**
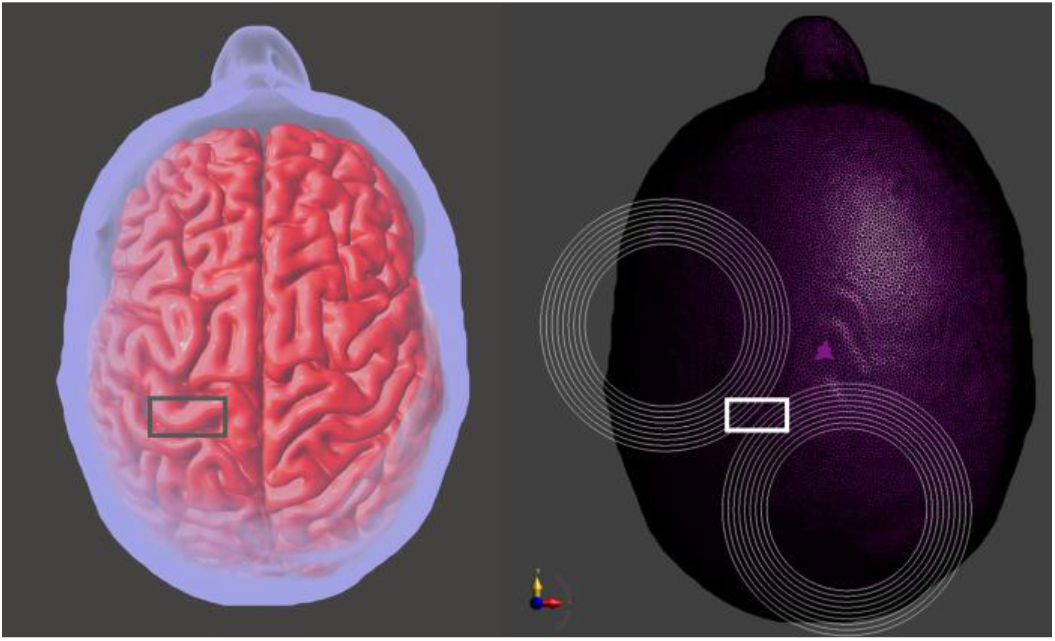
Fixed coil position. Left: the position of the coil on the gray matter, centered on the primary motor cortex region. Right: the figure of 8 TMS coil position on the head model.

Magneto quasi-static low-frequency solver was used to calculate the induced E-field. The magneto-static vector potential is calculated using the Biot-Savart law in equation 1. The E-field and the vector potential of the magnetic field are decoupled and divided into the solenoid and irrotational E-field components in equation 2. For the solenoid and irrotational fields, we obtain equation 3. The magneto quasistatic equation is implemented From equations 2 and 3 as shown in equation 4 [24] [43].

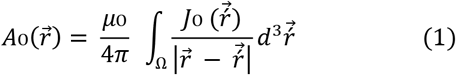

Where *A*_O_ is the magneto-static vector potential, *J*_O_ is the current density field, *μ*_O_ is the vacuum magnetic permeability, and d is the longest diagonal of the computational domain.

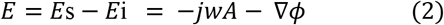

Where *E* is the E-field, *E*s is the solenoidal E-field, *E*_i_ is the irrotational E-field, *ϕ* is the scalar potential, w is the angular frequency and j is a complex number.

When:

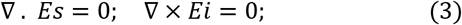

The magneto quasistatic equation becomes:

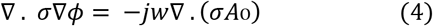

Where *σ* is the electric conductivity of the material.

Simulation results were verified to ensure the correct brain region was stimulated. Using slice viewer in Sim4life, a coronal slice of the brain was extracted from each head model representing the E-field profile shown in Fig. 4. Anatomical coronal slices were then taken at the same coordinate as the E-Field coronal slice to introduce a matching pair for training the DCNN. The maximum electric field intensity scale was set to a fixed 300 V/m scale to ensure all E-Field intensities are included in the study, in addition, to ensuring that all the input data sets are on the same scale.

**Fig. 4.**
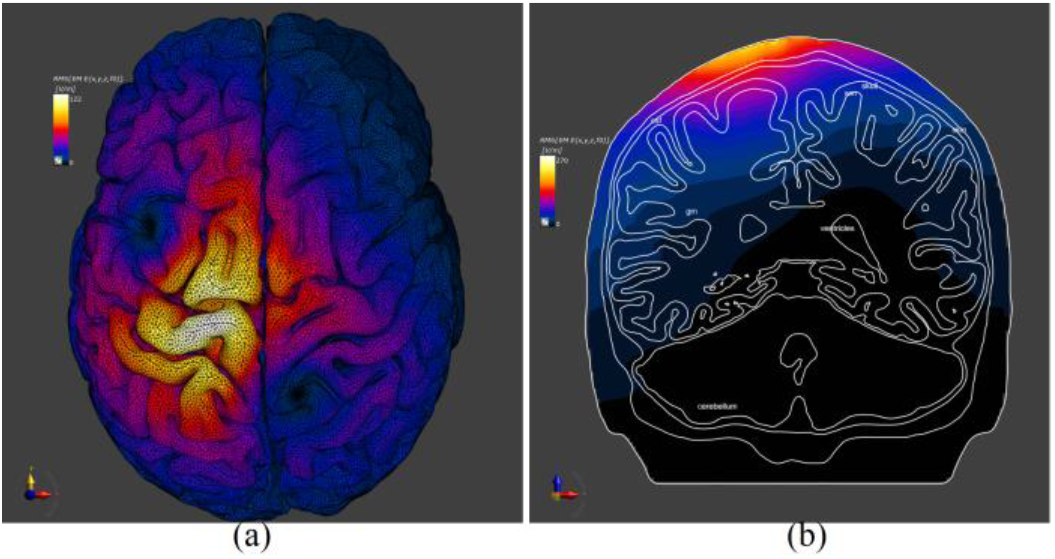
(a). The simulated induced E-field distribution on the gray matter for a representative subject. Maximum stimulation was located on the primary motor cortex. The maximum induced E-field was 122 V/m which is represented in white, wheras dark blue represents 0 E-field. Fig. 4 (b). Coronal slice view of the induced E-field on the head model. This slice has the maximum induced E-field in the x-z plane.

### E. Data Preprocessing

We first implement max-min normalization techniques to rescale the data into standard inputs for neural networks. Then we pad the images with zeros at the boundaries to make the image dimensions easier for downsampling [44]. Next, we split the data into training, validation, and testing sets randomly. The validation set is used to reduce the chances of overfitting.

### F. Data Augmentation

Due to the limited data generated from the head model, data augmentation [45] is implemented to generate ‘new’ training samples from the original ones by applying random jitters and perturbations. In addition, the proposed model can learn more robust features with more data to increase the generalizability of the model. We obtain the augmented data from the original MRIs generated from the head model by applying geometric transforms. Then, we pre-process the data with max-min normalization and zero-padding. Anatomical images and corresponding E-field data have been collected by us in a detailed data acquisition process (using SimNIBs pipeline and Sim4life); however, processing a large number of MRIs was not easy and it was time-consuming. Synthetic data augmentation techniques can be a probable solution to this problem. In our case, the data are collected from healthy subjects where TMS is applied to a specific region of the brain (primary motor cortex) for a fixed coil position. Therefore, considering these special aspects of data, augmenting solely based on rotation, flipping, mirroring, etc. on the same dataset does not make any significant change to predicted E-fields. Hence, we added the data augmentation process to the supplementary section.

### G. Model

A DCNN_based encoder-decoder network (U-net) is developed (Fig. 5) for the prediction of induced E-fields from MRI-based anatomical slices. We used 2D anatomical image slices with resolution (1024×1024) and E-fields with the same resolution as the data for training and testing the neural network model (Fig. 5). The encoder-decoder DCNN model can be trained with 3D images; however, it is unnecessary since the 2D slices we are using contain abundant contextual data. With the DCNN model, anatomical slices can be translated to 2D E-fields after optimizing the network parameters. Therefore, the network requires training at first, and then the trained network can predict corresponding E-fields from a new set of anatomical images. With this scheme, once the model is trained, it is possible to predict E-fields in real time, whereas FEM-based methods require hours to predict the E-fields.

**Fig. 5.**
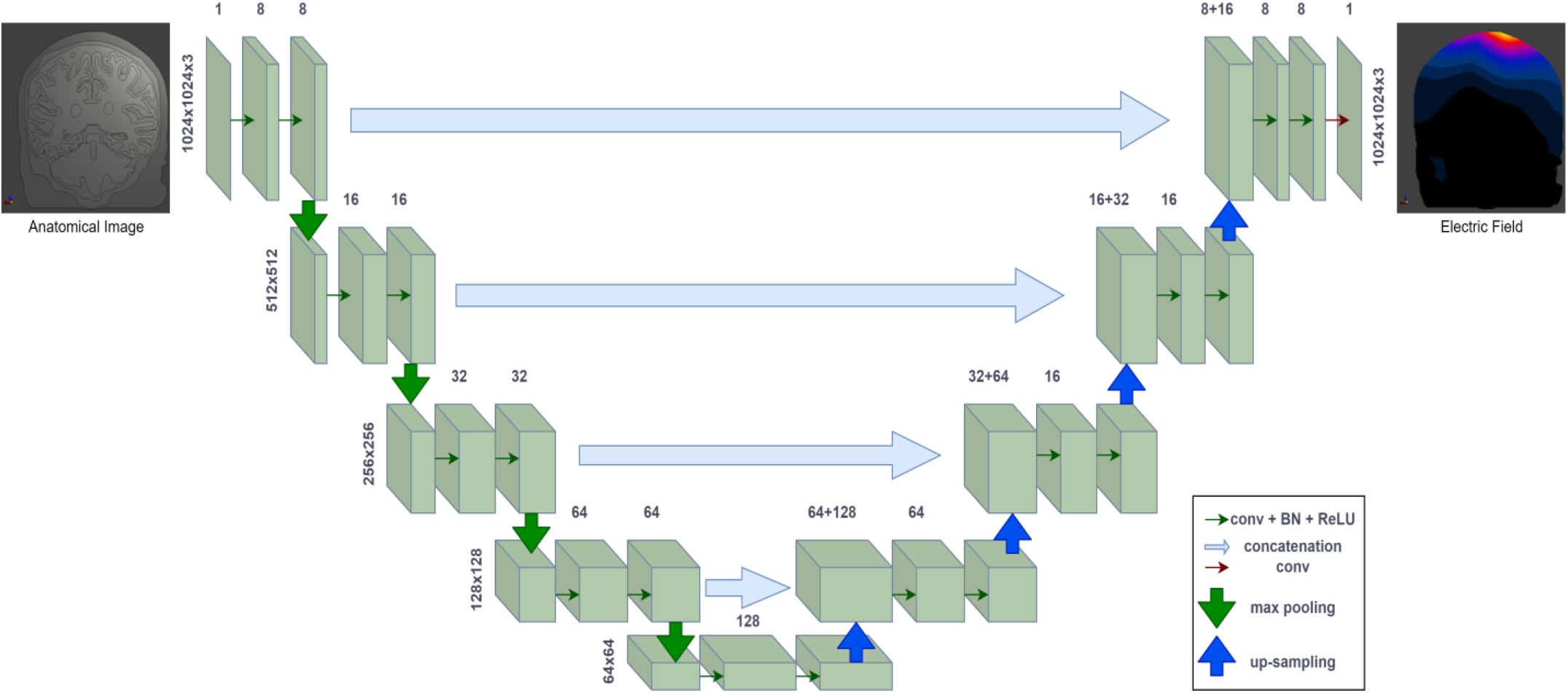
DCNN based encoder-decoder architecture used for prediction of E-fields using MRI based anatomical images. The horizontal green arrows represent convolution operation followed by batch normalization and a ReLU activation function. Vertical green arrows indicate max-pooling operations. Vertical blue arrows indicate up-sampling operation. The long horizontal light blue arrows denote concatenation/copying layers. Image size and number of channels for each convolution layer are provided along green bars. The network starts with input anatomical image data and ends with predicted output E-field.

As observed in Fig. 5, the DCNN model can be divided into two primary components: an encoder (left half) and a decoder (right half).

The encoder operates similarly to conventional convolutional neural networks, which can be trained to extract increasingly complicated relevant features from an image data. The encoder part contains multiple convolutions and max pooling layers. With a set of kernels, each convolution layer in the encoder performs a series of convolution operations on its input and then sends the outputs through a batch normalization (BN) layer and a nonlinear activation (ReLU) layer. Max pooling operation reduces the spatial dimension of the output image by keeping the most significant features. Mathematically, the operations in the encoder can be expressed as:

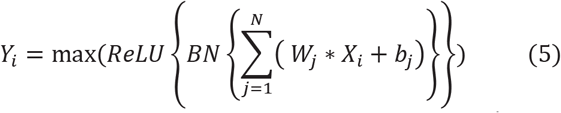

Here, *W*_*j*_ and *b*_*j*_ denote weights and bias of the j^th^ kernel, respectively. The asterisk symbol represents convolution operation. N denotes the total number of kernels in the i^th^ layer. *X*_*i*_ represents the input to the i^th^ layer and *Y*_*i*_ represents the output of the i^th^ layer. Anatomical image is used as the input for the first convolution layer. The successive convolution layers use the preceding layers’ output as input. For the network, four convolution layers are designed, and the number of kernels are 8, 16, 32, and 64, respectively, from the first to the fourth convolution layer with a kernel size of 3×3. Kernel weights are tensors with the same number of values as the input channels. Kernel weights and biases are free parameters that are updated in each iteration of the training to optimize their values. To make sure that input and output in each layer have same size, padding is used during the convolution process. The max pooling is performed with a size of 2×2 and stride 2 (non-overlapping). Hence, it compresses the spatial dimension of the extracted features by half. The pooling layer has no free parameters that can be learned. However, for later use in the decoder, the locations in each pooling window where the highest value is located must be preserved.

The decoder is designed to be a mirrored edition of the encoder where a gradual reconstruction of the E-field is done with an additional convolution layer added at the end to transform each feature vector from the preceding layer to the predicted E-field. The spatial dimension of the predicted E-field is identical to the input anatomical image (1024×1024), as shown in Fig. 6.

**Fig. 6.**
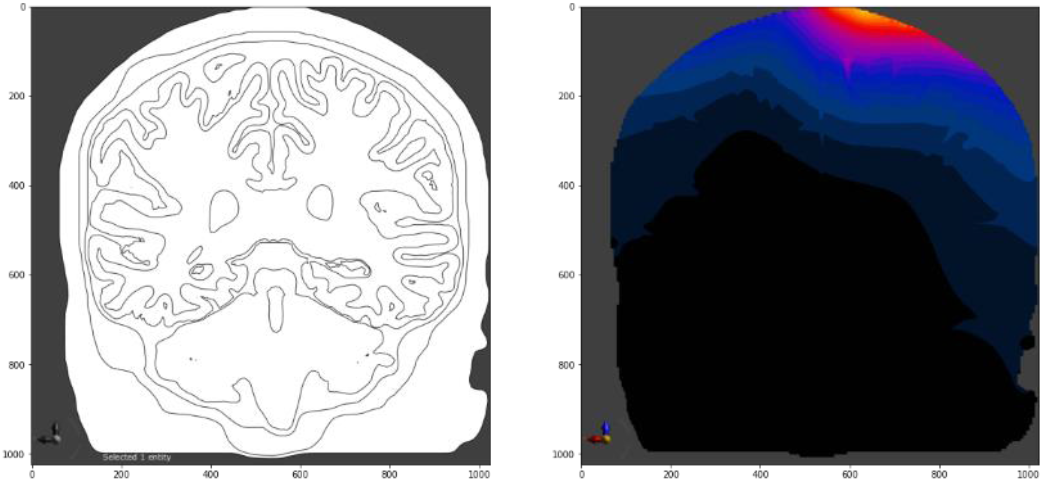
A sample pair of anatomical MRI and corresponding E-field used for training the learning network.

The decoder uses multiple up-sampling layers to transform data from dense to sparse resolution, in contrast to the feature extraction done in the max pooling layers, where gradual reduction of the spatial dimension of the features is observed. Each up-sampling operation doubles the input features’ spatial resolution. To allow the extracted complex features from the encoder to be used as additional inputs for the convolutional layers in the decoder, skip connections [27] from the encoder to the corresponding decoder are implemented (shown as horizontal light blue arrows in Fig. 5. Skip connections copy the complex features from the encoder and combine them with upsampling layer’s outputs in the decoder using preserved location information for the pooling windows in the encoder. High dimensional predictions are performed more smoothly in the decoder because of these skip connections. Convolution operations are then performed in the decoder convolution layers to convert the sparse features into dense features. The convolution layers in the decoder have their own set of learnable weights and biases that are uncorrelated to the corresponding encoder weights and biases.

Training the encoder-decoder DCNN model requires estimation of all learnable network parameters, which can be accomplished by minimizing the prediction error (loss function) between predicted E-fields and corresponding ground truth E-fields. Considering sample anatomical images as {*X*_*i*_} and their corresponding reference E-fields as{*Y*_*i*_}, the mean squared error (MSE) loss function can be expressed as:

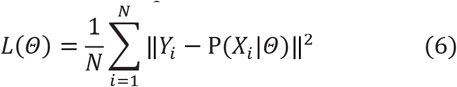

Where *Θ* is a set of learnable model parameters and P(*X*_*i*_|*Θ*) is predicted E-fields for a sample image *X*_*i*_. N denotes the total number of images.

The data used for training the encoder-decoder network consisted of 187 anatomical images and their corresponding E-fields, which we split into 168 pair (training) and 19 pair (validation) datasets. A total of 200 iterations were used for training the network parameters. Each iteration is known as an epoch of the training.

## III. Results

Simulation results were verified by ensuring the E-field hotspot magnitude was located on the motor region (M1) of the gray matter segment [42]. Scaled head models for the same subject presented different y-coordinates for the maximum E-field intensity slice in the x-z planes. This variation of maximum E-field intensity slice coordinates indicates that scaled head models represent distinct brains, which allowed us to consider them as more data inputs for machine learning algorithms. The first two columns in table I show 11 subjects’ maximum E-Field coronal slice coordinates variations, which impose inter-subject variability between the head models. Columns 3 and 4 show maximum E-Field coronal slice variation for the head model of subject 3 after scaling.

**Table I:** Maximum E-Field Coronal Slice Coordinates

| Subject | Max E-Field Y-coordinates in (mm) | Subject 3 Scales | Max. E-Field Y-coordinates in (mm) |
| --- | --- | --- | --- |
| 1 | -39.78 | 0.96 | -34.11 |
| 2 | -35.28 | 0.97 | -31.69 |
| 3 | -32.45 | 0.98 | -32.85 |
| 4 | -23.51 | 0.99 | -32.21 |
| 5 | -33.54 | 1 (original scale) | -32.45 |
| 6 | -29.73 | 1.01 | -33.65 |
| 7 | -23.16 | 1.02 | -34.41 |
| 8 | -35.33 | 1.03 | -35.27 |
| 9 | -26.63 | 1.04 | -34.14 |
| 10 | -20.09 | 1.05 | -31.46 |
| 11 | -21.13 | 1.06 | -31.56 |

Training can be a computationally intensive process. However, adding enough GPU memory will accelerate the process. The model training with 200 epochs takes around two hours with added GPU as hardware accelerator in the Google Colaboratory. After the model is trained, test data prediction can be calculated in real-time since only inference is required.

Fig. 7 illustrates training and validation loss (MSE) for each iteration of the training network. Training loss decays exponentially over iterations. Validation loss in Fig 7, however, shows some random fluctuation over the iterations and settles down at the end of training. Fig. 8 shows a sample of three representative E-field results as produced by the proposed DCNN method, along with the corresponding reference or ground truth E-field. For our trained network, MSE for training and validation are 5.215×10^−4^ and 1.818×10^−3^, respectively. Training and validation data peak signal to noise ratio (PSNR) are 32.83dB and 27.4dB, respectively. Therefore, the results show good accuracy of prediction. For most parts of the head region, the DCNN method predicts reasonably accurate values as seen in Fig. 8. Large errors mainly occur at interfaces between different tissue types, especially around the borders.

**Fig. 7.**
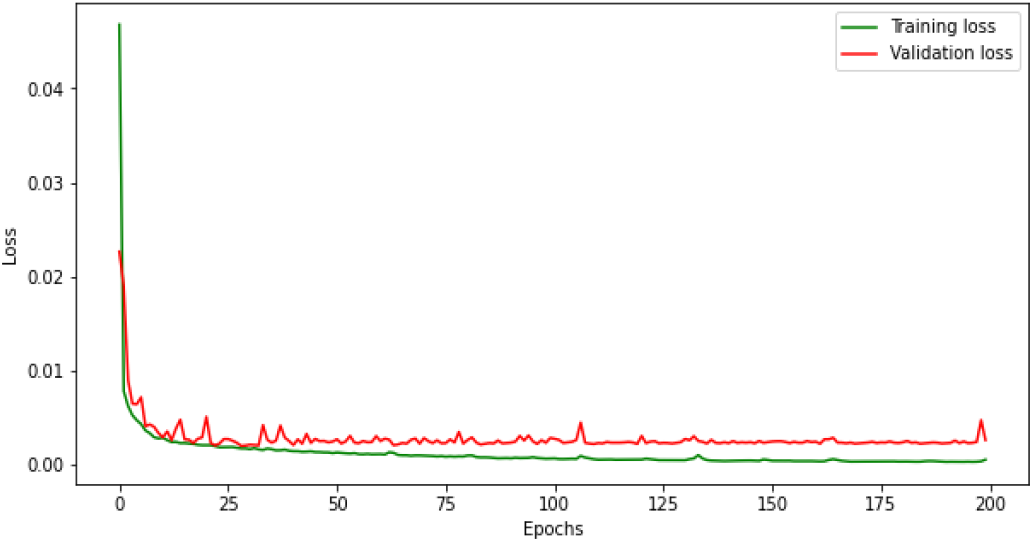
Training and Validation loss in each iteration of training. Training loss decays exponentially over iterations. Validation loss however shows some random fluctuation over the iterations.

**Fig.8.**
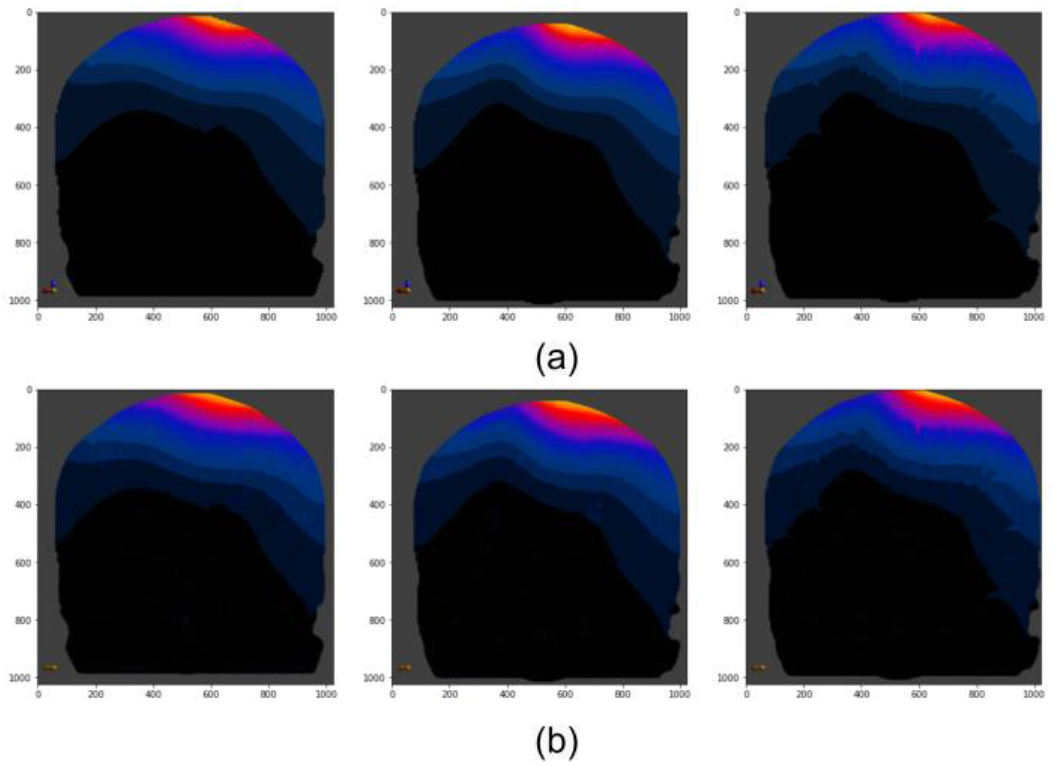
Comparing samples of (a) ground truth E-fields, (b) predicted E-fields.

This is due to high-intensity gradients in these areas, but it may also be partially due to the non-perfect alignment between the anatomical images and ground truth E-fields.

## IV. Discussion

In this study, we acquired accurate prediction of the E-fields in the desired area. This prediction could be improved by including more images for the same head model; for instance, increasing the number of subjects’ MRIs instead of scaling the original head models will embrace more differences concerning inter-subject variability, which will include more head model variations for training the algorithm. 3-Dimensional E-Field and anatomical slice images could be used as input for machine learning algorithms to include more brain anatomy in the training algorithms; however, it increases the computation resource and time significantly. Additionally, including fiber tracts in the head model and finite element simulations will add more accuracy to the study and create a close dependency between the threshold E-field and RMT.

The results of the model after data augmentation compared with the original model show that the model achieves a similar mean square error and peak signal-to-noise ratio after using MRI images with data augmentation. However, after training with more instances, the validation loss becomes more stable.

We conjecture that the reason for getting similar results is that the data augmentation methods are simple. The newly generated data contains some noise which may reduce the model performance. For data augmentation, we only considered rotation, shifting, and zooming. However, these techniques do not introduce new synthetic data to the model. Instead, we only include the same samples in a different state, resulting in a limited impact on generalizability.

Besides these basic data augmentation techniques such as rotation, shifting, and zooming, we could also consider other techniques such as cropping, brightening, adding Gaussian noise, and so on. In addition, we also consider deep learning-based methods to augment existing limited data. For example, conditional GANs [46] can be used to transform an image from one domain to an image to another domain. GANs take random noise from a latent space and produce unique images that mimic the feature distribution of the original dataset. Other modified versions of GANs, such as self-supervised GAN [47], CycleGAN [48] can also be used to generate synthetic images to augment the training data.

The E-field modeling carried out by most of the researchers in the field consider each layer of the brain with homogenous electrical properties, which doesn’t accurately represent the brain. In our ongoing project, we have obtained fiber tract images through diffusion tensor imaging (DTI) and each subject’s RMT. We are working on integrating fiber tracts in E-field modeling for an accurate representation of brain geometry and heterogeneous electrical properties, which is outside the scope of this work. We plan to use the accurate E-field modeling data for training in DCNN to predict the E-field that may correlate well with the RMT obtained from the 11 subjects recruited in this project.

## V. Conclusion

The DCNN model predicted corresponding E-fields in real-time with training and validation PSNR of 32.83dB and 27.4dB, respectively; thus, DCNN can be an alternative to time-consuming methods using FEM alone.

In this study, one anatomical-coronal slice for each head model scale was used as an input for the machine learning algorithm to get the predicted maximum induced electric field on the targeted regions. The model predicts E-fields based on anatomical variations in the brain. However, this prediction can be used for just one type of TMS coil. Depending on the material and geometry of the TMS coil, induced E-fields will be different, and each TMS coil setting will require separate data for constructing and training a neural network model for estimation of stimulation strength and induced E-fields. The difference in learning models and parameters based on coil variation is yet to be solved.

## Acknowledgment

Authors would like to thank Dr. Neil Mittal for recruiting the subjects in this study and Dr. Cooper Hodges for acquiring the subjects’ MRI’s.

