## Supplementary Material for "Prediction of Electric Fields Induced by Transcranial Magnetic Stimulation in the Brain using a Deep Encoder-Decoder Convolutional Neural Network"

**Data Augmentation:**

**Methods**: We first expand our dataset via data augmentation. We randomly selected 25 gray-scale MRI images and implemented data augmentation to generate a new dataset with 231 MRI images. Then, we initialize the ImageDataGenerator object to perform random translations, rotations, zooming and shifting on our input images. The generator saves these augmented images as ".png" files to the specific output directory as the new dataset. Finally, we loop over these augmented images from our image data generator and count them until reaching the final required number of augmented images. The parameters for data augmentation are: rotation range: 90, width shift range: 0.1 height shift range: 0.1, and zoom range: 0.2

Rotation: Image rotation is a commonly used augmentation technique that allows it to become invariant to the orientation of the object. We randomly rotate the image from 0 to 90 degrees. When the MRI image is rotated, we replace the empty areas with the nearest pixel values when the pixels move outside the image. Shifting: We shift the pixels of the MRI images horizontally and vertically by adding a constant value in the range of 0.1 to all the pixels. Zooming: we implement both zoom-in and zoom-out on the MRI images with the range of 0.8 to 1.2.

**Experiment:** We first used the MRI data consisting of only 12 (training) and 2 (validation) images with a batch size of 1 for training. We obtain Mean Square Error (MSE) as 0.0048 and Peak Signal to Noise Ratio (PSNR) as 23.19dB. Then we apply data augmentation as discussed in Section E to generate more training data. After rotating, shifting, and flipping the original images, we got 140 MRI and E-field images, including the 14 original data and 126 generated data. We randomly split the data into training and validation sets with 9:1, respectively. After training the model with the augmented images. Fig. 2 shows the predicted E-field in comparison with the ground truth. We achieved MSE as 0.0027 and PSNR as 25.68dB for the validation set. Figure 2 demonstrates the comparison of training loss and validation loss for the data augmented DCNN.

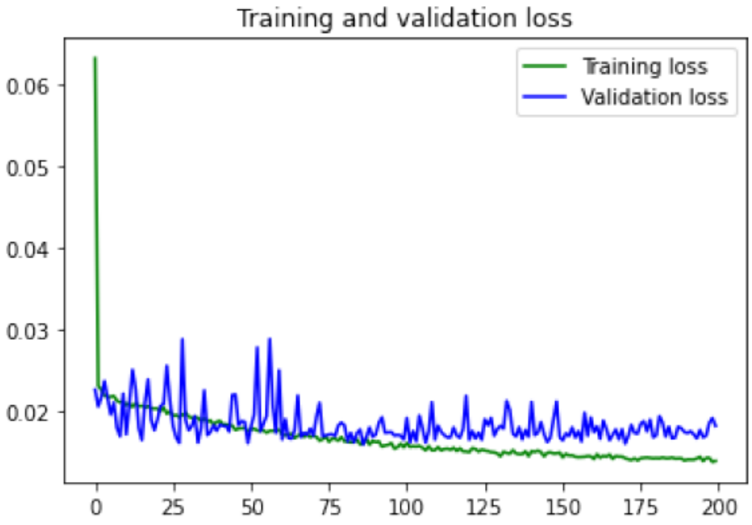


Fig. 1. Model Training and validation loss after data augmentation. MSE: 0.018262804, PSNR: 17.38dB

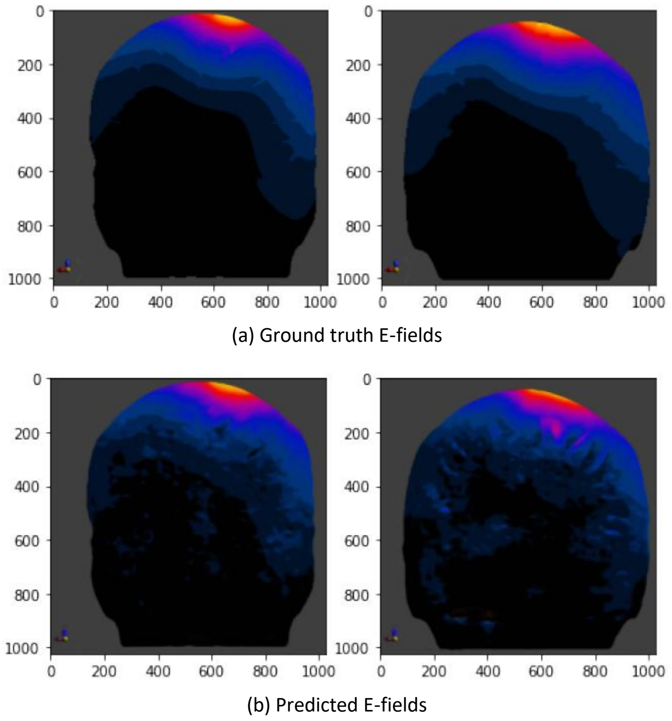


Fig. 2. Comparing samples of ground truth E-fields (a) to predicted E-fields (b).
